# Closely Related Tree Species with Overlapping Ranges Exhibit Divergent Adaptation to Climate

**DOI:** 10.1101/2021.11.08.467758

**Authors:** John W. Whale, Collin W. Ahrens, David T. Tissue, Paul D. Rymer

## Abstract

With global climate change shifting and altering temperature and precipitation regimes, the ability of natural forest stands to persist in their local environments are being challenged. For many taxa, particularly among long lived tree species, the potential to respond is underpinned by genetic and trait diversity and may be limited. We sampled 326 and 366 individuals of two widely distributed and closely-related red gum *Eucalyptus* species (*E. blakelyi* and *E. tereticornis*) from across their entire Australian range. We identified putatively adaptive variants associated within genes of key biological processes for both species. We mapped the change of allele frequencies of two hierarchical gene ontology groups shared by both species across geography and climate and predict genomically vulnerable regions under a projected 2070 climate scenario. Regions of potential vulnerability to decline under future climate differed between species and may be applied to guide conservation and restoration strategies. Our study indicated that some populations may contain the adaptive genomic variation necessary for these species to persist through climate change, while others may benefit from the adaptive variation of those populations to enhance resilience.

## INTRODUCTION

Local adaptation provides higher fitness of local populations based on environmental conditions at their home site (Savolainen et al., 2007, 2013) resulting from natural selection over many generations. The rapid rate of climate change is putting pressure on natural populations to adapt in order to persist, particularly among long-lived and sessile organisms such as trees (Kawecki & Ebert, 2004). Species within the greatest adaptive genomic variation will be best placed to persist with changing climate (Alberto et al., 2013). However, should species, or locally adapted populations fail to respond to these environmental pressures, they may still be at risk of local decline or extinction (Aitken et al., 2008). Therefore, it is critical to determine the distribution of adaptive genomic variation in forest species to improve predictions of vulnerability and management strategies going forward.

Species adaptation can manifest through several pathways; standing genomic variation throughout the genome, via new mutations, the introduction of adaptive variants through hybridisation and introgression (Bragg et al., 2015), or whole genome duplication and polyploidy (see review by Hollister, 2015). Variants may be genic (located within genes or coding regions), regulatory (located near genes), or non-genic (outside gene regions, and may be referred to as ‘neutral’). It is the variants in genic or regulatory regions that alter the phenotype (amino acid sequence or expression of a protein; Weigel & Nordborg, 2015; Zou et al., 2017) and therefore may be subject to selection pressures. Through the characterisation of genomic variants and the genes with which they interact may provide useful insights into the adaptation of species to their environment and climate.

Temperature and precipitation are known to limit species’ distributions, while also influencing adaptation (Blackman et al., 2014; Savolainen et al., 2007). With temperature and rainfall regimes projected to change in the future, species may become maladapted at local scales since beneficial alleles may be at low frequencies or locally non-existent, potentially leaving some species playing catch-up. Fortunately, genotype-environment association (GEA) tools can be employed to elucidate signatures or patterns of local adaptation throughout a climate landscape. These methods are being increasingly utilised on a variety of model and non-model organisms, including plants and forest trees (Ahrens et al., 2018). They may be employed on large genomic datasets to identify alleles correlated with an environmental or phenotypic variable. Recent studies have utilised these methods to detect candidate variants and allelic shifts across the landscape inferring local adaptation in *Populus* (Evans et al., 2014; Fahrenkrog et al., 2017), *Quercus lobata* (Gugger et al., 2021), and among eucalypts (Ahrens et al., 2019; Jordan et al., 2017). Through the characterisation of candidate variants, researchers may begin to predict the genetic risk posed on species as a result of climate change (Breed et al., 2019; Browne et al., 2019). Such data may then be presented to land managers for decisions into effective strategies for long-term observation and/or translocations.

Australia’s widespread red gum eucalypt species (family Myrtaceae, section *Exsertaria*), here specifically *E. blakelyi* Maiden and *E. tereticornis* Sm., are of social, cultural, economic, and ecological importance. Forests of these species have provided indigenous communities with medicinal resources and wood for building canoes, hunting tools and ceremonial paraphernalia (Australian National Botanic Gardens, 2004). Economic benefits are recognised both nationally and internationally where natural and planted forests may be used in forestry for building materials in the housing or transport industries (Ginwal et al., 2004; Kaur et al., 2011). They are of huge ecological importance through providing key ecosystem services (de Carvalho Balieiro et al., 2020) and habitats for many native fauna. Further, these trees are dominant upper-storey species within the critically endangered Cumberland Plain woodland (New South Wales Department of Planning, Industry and Environment, 2019a) and the threatened White Box-Yellow Box-Blakely red gum (New South Wales Department of Planning, Industry and Environment, 2019b) ecological communities in both remnant and fragmented populations (Gibbons & Boak, 2000). Additionally, populations of *E. blakelyi* have suffered from regular dieback particularly around the Australian Capital Territory (ACT) with reports dating back some near-40 years (Landsberg, 1985, 1990; Landsberg & Wylie, 1983). Both red gum species are latitudinally and longitudinally widespread s, which makes this group prime candidates to study the adaptation to climate of *Eucalyptus* species since they traverse several climatic gradients.

Here, we aimed to better understand the underlying adaptive genomic variation among widespread *Eucalyptus* species associated with their climatic origin, and also provide a comparative framework of two closely related species across climatic gradients. The species distributions are partly overlapping along eastern Australia, however *E. blakelyi has* greater rainfall variation distributed on the western slopes and *E. tereticornis* occurs largely east of the Great Dividing Range with greater temperature range. We surveyed the genomic variants (single nucleotide polymorphisms; SNPs) of natural populations with contrasting distributions and climate origins to improve our understanding of the genomics of local adaptation and adaptive capacity to climate change. Testing the following hypotheses (i) both species will show significant genomic signatures of adaptation to climate; (ii) putatively adaptive variants associated with temperature will explain more of the total variation compared in *E. tereticornis* compared to *E. blakelyi*, while variants associated with precipitation explains more variation in *E. blakelyi*; (iii) Gene functions underlying the adaptive genomic variation will predict local adaptation to climate, and (iv) populations currently experiencing the extremes in temperature and rainfall variables will be most at risk of decline under future climate scenarios.

## METHODS

### Study Species, Site Selection and Sampling

In this study, we analyse two widely distributed red gum *Eucalyptus* species endemic to eastern Australia, *Eucalyptus blakelyi* Maiden (Blakely’s red gum) and *E. tereticornis* subsp. *tereticornis* Sm. (forest red gum). Each species traverses variable climatic gradients across eastern Australia. Both species are key foundation species in the grassy woodland ecosystems they inhabit and form tall trees reaching 30m in *E. blakelyi* and 50m in *E. tereticornis*. However, both species’ distributions are largely separated by the Great Dividing Range along Australia’s east coast. Further, species may be distinguished by bud morphology where they are both longer than they are broad (more than 2.5x so in *E. tereticornis*), and on occasion be pruinose in *E. blakelyi* (Brooker & Slee, 2000; Klaphake, 2012).

For both species, sampling locations were identified based on current occurrence records downloaded from Atlas of Living Australia (ALA; ala.org.au) to cover their geographic and climatic distributions. These records were filtered to remove observations beyond the natural ranges of each species (e.g. botanic gardens and urban plantings), ambiguous species identification, and observations made prior to 1990 given the poor spatial locality information of old records and changes in land-use decreasing the likelihood of sampling. The remaining data were analysed through pairwise bioclimatic variation (Fick & Hijmans, 2017) downloaded at 30-second resolution (∼1Km per pixel). Potential populations were shortlisted based upon the similarity and dissimilarity of these pairwise climatic variables based on geographical distance (i.e. similar climate space among distal populations, and dissimilar climate space among proximal populations; Figure S1), and were further filtered to remove, where possible, populations <10 Km distance and inaccessible sites (using Google Maps Street View).

From throughout their natural distributions, a total of 326 mature trees for *Eucalyptus blakelyi* from 34 sites spanning 1,096 Km (2014 range size 509,075 Km^2^), and 366 individuals for *E. tereticornis* subsp. *tereticornis* (hereon referred to as *E*. tereticornis) from 38 populations spanning 2,381 Km were sampled (2014 range size: 792,575 Km^2^; González-Orozco et al., 2016) (Figure 1). Leaf material was collected from 6-11 individuals per species per site, with individuals at least 20 metres apart. Leaves were kept cool and dark before later freeze-drying and subsequently kept dry on silica gel in a container at room temperature.

**Figure 1:**
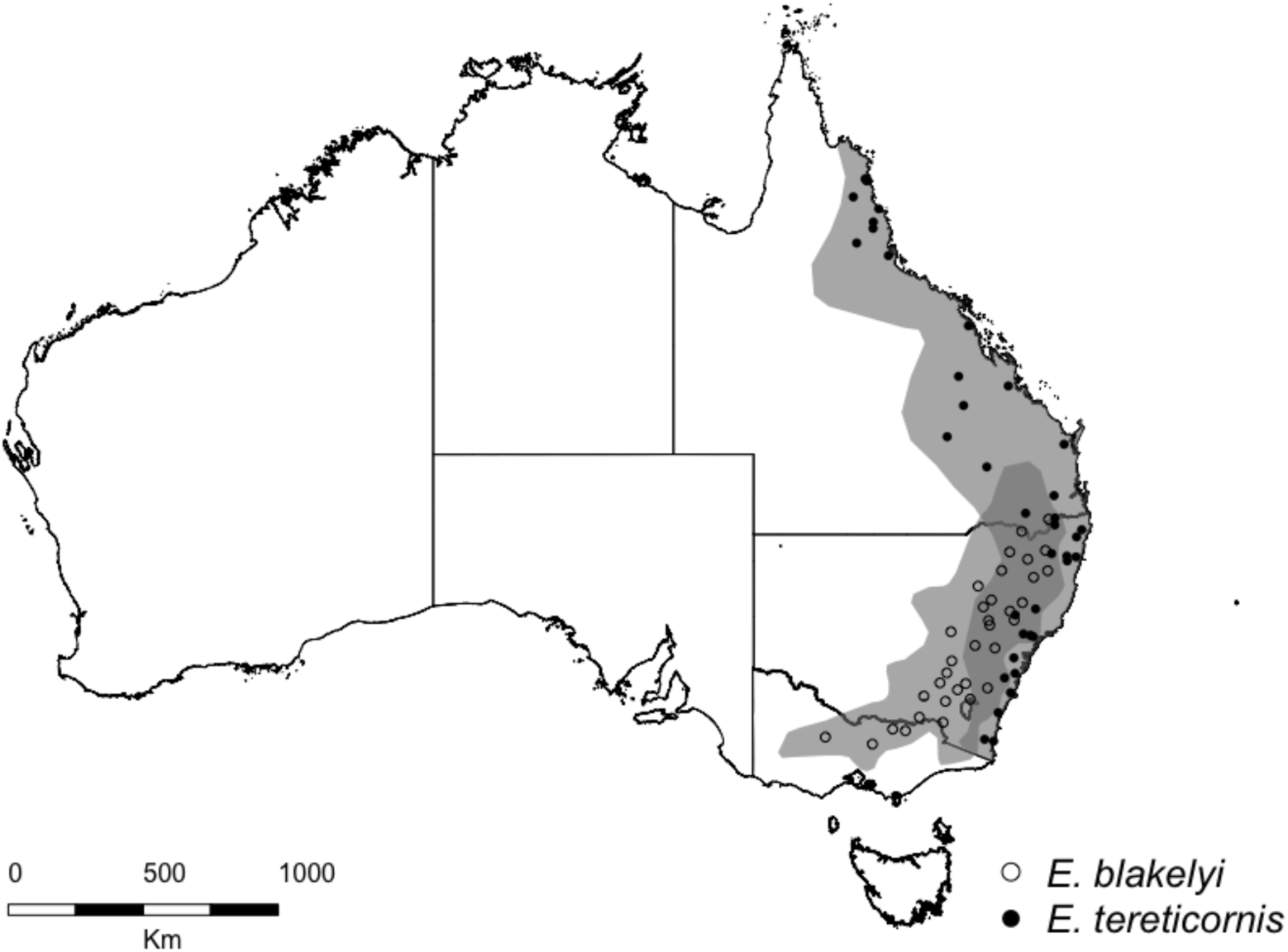
Species distributions (grey) and sample sites for *E. blakelyi* and *E. tereticornis*.

### SNP Generation and Bioinformatics

We sent 10-15 mg of freeze-dried leaf material from each individual to Diversity Arrays Technology Pty Ltd. (Canberra, ACT, Australia). DNA was extracted using a modified CTAB protocol and samples sequenced using the DArTseq protocol; a reduced-genome complexity next-generation sequencing method (Kilian et al., 2012; Sansaloni et al., 2011). Reduction of the genome is conducted through the use of a combination of *Pst*I and *Hpa*II restriction enzymes with fragments cleaved by both enzymes ligated with sequencing adapters. Sequences were filtered with a minimum quality threshold phred score of 30. SNP filtering was performed separately for both species in R (R Core Team, 2019) using the DARTR package (Gruber et al., 2018). SNPs were retained if called in 223 individuals and 249 individuals respectively (35% missing data). This threshold was chosen due the high numbers of individuals among both species, and as such would still provide high power for performing population-level allele frequencies and genotype-environment association (GEA) analyses (Ahrens et al., 2021; Lotterhos & Whitlock, 2015). Minor allele frequency (MAF) was set to 0.02 – that an allele call would need to be present in at least two populations to be retained due to the total number of populations and individuals genotyped. Further, one SNP scored per 69 bp sequence read was randomly retained. Linkage disequilibrium (LD) was calculated via the *snpgdsLDMat* function and pruned at 0.5 similarity using the *snpgdsLDpruning* function in SNPRELATE (Zheng et al., 2012).

### Population Differentiation and Genotype-Environment Association Analyses

Population differentiation for both species was estimated using the Weir and Cockerham *F*_ST_ method (Weir & Cockerham, 1984) in the HIERFSTAT package (de Meeûs & Goudet, 2007; Goudet, 2005). To investigate population structure further, we employed a discriminate analysis of principal components (DAPC) analysis (Jombart et al., 2010), a multivariate approach that identifies and defines clusters of related individuals. We first used the *find*.*clusters* function to determine the number of clusters within each dataset in the ADEGENET package (Jombart, 2008; Jombart & Ahmed, 2011). The *find*.*clusters* function partitions between- and within-group variance through an increasing *k*-means approach (Jombart et al., 2010). The number of sub-population (or clusters) is determined by the lowest Bayesian Information Criterion (BIC) score. The *dapc* function allows for visualisation of the population structure without *a priori* knowledge of the number of the clusters within the dataset by first transforming the data using principal component analysis (PCA) followed by characterisation of clusters using discriminant analysis. Here we retained 230 principal components for *E. blakelyi* and 225 for *E. tereticornis* respectively, with 2 discriminant functions retained for both species and visualised on a 2D scatter plot.

Prior to testing for the associations between climate and SNPs, we determined the spatial dependence of environmental variables to one another within the sampling distribution for each species. A Pearson’s correlation coefficient (*r*) was calculated for 20 climatic variables (19 bioclimatic variables plus annual aridity index) using the *corr* function in the VEGAN package (Oksanen et al., 2019). We ensured that climates used in one species were used in the other to test hypotheses about divergent evolution. In total, eight climate variables were chosen based on *a priori* knowledge of the species and spatial independence from the PCA. Additionally, a *Moran’s I* was calculated to determine the spatial autocorrelation of eight climatic variables within both species (BIO1; mean annual temperature (T_MA_), BIO4; mean temperature seasonality (T_SN_), BIO5; mean maximum temperature of the warmest month (T_Max_), BIO6; mean minimum temperature of the coldest month (T_Min_), BIO12; mean annual precipitation (P_MA_), BIO14; mean precipitation of the driest month (P_DM_), BIO18; mean precipitation of the warmest quarter (P_WARMQ_), and annual aridity index (AAI)) (see suppl. Table S1 & S2) using the *Moran*.*I* function in the package APE (Paradis & Schliep, 2019) to determine the effective sample size (ESS) of each population based on environmental independence. It is worth noting that the eight climatic variables used on both species were not entirely independent, but showed low correlations within one species or the other, though values of *r* up to 0.7 do not increase the number of false positives detected (Ahrens et al., 2021).

To test the hypotheses of climate-associated genomic variants of these two eucalypts, GEA analyses were performed to detect SNPs putatively under selection (hereon referred to as ‘putatively adaptive SNPs’). We used three methods that use different models to detect SNP-environment correlations. A multivariate method, redundancy analysis (RDA), was performed to understand the relationship between genomic variants and climate. As RDA requires a full dataset, missing data values were imputed, with the RDA analysis performed in the VEGAN package 2.5-6 in R using the *rda* function. We used the *anova*.*cc* function to estimate the overall significance of the analysis. If significant, we then tested the significance of each significant climate variable using 999 permutations in the significance tests. In addition, two univariate methods, BayPass and LFMM2, were employed. BayPass v2.2 (Gautier, 2015), a Bayesian method which builds on the algorithm used in BayEnv and are discussed in Gautier (Gautier, 2015), which utilises population-level allele frequencies to determine associations between climate and SNPs, and controls for population structure by employing an Ω (*F*_ST_analog) covariation matrix (Günther & Coop, 2013). We ran the core model five times on each dataset using 20 pilot runs with 500 iterations, 5,000 Burn-in, and 1,000 iterations, to acquire five independent Ω matrices. A mean of these matrices was calculated in R, that was used as the covariance matrix in the auxiliary (AUX) model for determining genomic associations to climate. Covariates (climate data) were scaled using the *scalecov* function and the MCMC run with the *auxmodel* function for the auxiliary covariate model to estimate the Bayes Factor (BF). The parameters for the AUX model were also 20 pilot runs with 500 iterations, 5,000 Burn-in, and 1,000 MCMC iterations. Associations were considered when the BF significance threshold was ≥ 20 (Kass & Raftery, 1995).

We used LFMM2 (Caye et al., 2019), a new algorithm that builds on the previous LFMM (latent factor mixed model) program using an exact solution of a regular least-squares problem, to identify genetic associations to climate. Here, we inputted individual genotypes, and imputed missing data using the *impute* function with the median method used to complete the genotype-data matrix. LFMM2 accounts for population structure by using latent factors to estimate the discrete number of ancestral clusters (*K*) contributing to the genetic variation via a PCA. Using this estimate for each species, we used the *lfmm_ridge* function to fit the lfmm model of *K* factors. We then used the *lfmm_test* function to perform individual associations with each climate variable with the fitted model calibrated with the genomic inflation factor control method *gif*. Multiple testing can be done using Bonferroni corrections applied to the calibrated p-values. We applied a significance value of (α = 0.001 since lower thresholds were found to be more variable and returned greater numbers of false positives (Ahrens et al., 2021). These thresholds were applied to each of the eight climate variables.

### Characterisation of Putatively Adaptive SNPs

SNPs identified as putatively adaptive through the GEAs were aligned and annotated to the *E. grandis* genome (Myburg et al., 2014) using the blast function through the EucGenIE database (https://eucgenie.org, Christie *et al*., personal communication). Sequences with mean bit scores for chromosome-level alignments greater than 60.0 were retained using EUCANEXT (Nascimento et al., 2017). Chromosome and location of the candidates were recorded in relation to their alignment around coding regions (CDS) e.g. within coding region (CDS fragment recorded), putatively regulatory region to specific CDS fragment (within 1Kb of a CDS; Vandepoele et al., 2009), or non-genic (beyond 1Kb of a CDS). Identified sequences within coding or regulatory regions were analysed for their gene ontology (GO) term enrichment using the PlantRegMap online database (Tian et al., 2020). Enriched GO terms were examined for functions that were anticipated (e.g. response to stressors), and those common among species.

### Landscape Genomics Modelling

We performed a generalised dissimilarity model (GDM), a statistical approach to ascertain the importance of the climate variables upon our genomic SNP data and provide a visualisation of these spatial interactions across the landscape (Ferrier et al., 2007). We employed this method to characterise the allelic turnover of putatively adaptive SNPs identified from the GEAs, and of the loci identified under the common gene functions (GO term) found in both species. A pairwise genetic distance matrix was generated with the SNPs within the HIERFSTAT package using the *genet*.*dist* function. The distance matrices were created separately using the group of SNPs for each common GO term for each species. A generalised dissimilarity model within the GDM package (Fitzpatrick et al., 2020) using the *gdm* function was performed on the SNPs generating spline (deviance) plots through each climate variable. From these plots, only BIO18 and AAI for *E. blakelyi* for each GO term, and BIO1, BIO4, BIO6, BIO12, BIO14, and BIO18 were used for oxidoreductase activity, BIO1, BIO4, BIO12, BIO14, and BIO18 for tetrapyrrole binding, and BIO1, BIO4, BIO6, BIO12, BIO14, BIO18, and AAI for oxidation-reduction process for *E. tereticornis* were retained. To generate the projection of the GDM onto the current landscape, the occurrence records for both *E. blakelyi* and *E. tereticornis* were downloaded from the ALA and records beyond the natural species’ ranges were removed. A polygon of each species was created with the occurrence data using the free and open source QGIS v2.18. The deviance splines of the GDM models and stacked environmental rasters were transformed using the function *gdm*.*transform* and a PCA conducted of the transformed layers using *prcomp*. Spatial distribution of the layers were plotted using the first two principal components for *E. blakelyi*, and three components for *E. tereticornis* on a red-green-blue plot using the *ggRGB* function in RSTOOLBOX.

Further, we then used this model to predict spatial genomic vulnerability throughout the distributions of each species. For these predictions, we used the same seven bioclimatic variables used within the GDM models with the current climate data. Climate data predicted for 2070 were downloaded for the CMIP5 at 2.5 m arc spatial resolution based on the GCM CCSM4 and the 8.5 representative concentration pathway (RCP) (WorldClim, 2020) which is an emissions scenario of greenhouse gases, aerosols, and other climatic drivers. The 2070 climate data were also projected using the species’ distribution polygons. To predict the genomic vulnerability across the landscape, we used the current climate rasterstack data and model and subtracted it from the projected 2070 scenario rasterstack variables using the *predict* function and with *time = TRUE*, and the difference between the models plotted as above.

## RESULTS

### SNP Dataset and Population Structure

DArTseq returned a total of 70,793 SNPs, which resulted in 12,470 independent SNPs for *E. blakelyi*, and 12,466 for *E. tereticornis* that were distributed throughout the genome. On these full SNP data frames we estimated population differentiation across all samples within species, observing background levels of population structure for both species (*F*_ST_ = 0.089 (0.058-0.161) for *E. blakelyi*, and *F*_ST_ = 0.129 (0.056-0.185) for *E. tereticornis*). The number of clusters detected was K = 2 for both species, although *E. blakelyi* may possess more subpopulations since the curve began to rise more sharply after *K* = 5 (Figure S2). DAPC analysis conducted for *E. blakelyi* accounted for 83% of the variation, while for *E. tereticornis* represented 77.5% of the total variance (Figure S3) across the first two DA eigenvalue axes. Membership probabilities for each group for *E. blakelyi* were between 0.7 and 1, indicating low admixture and are distinct population groups. Meanwhile, for *E. tereticornis*, five of the 38 populations possessed probabilities outside this range (all from Queensland, between 0.3 and 0.5556), suggesting that these five populations experience greater admixture within each species.

### Genotype-Environment Associations

The GEA analyses identified a total of 328 SNPs for *E. blakelyi*, and 402 SNPs for *E. tereticornis* that were found to be significantly associated with at least one climate variable. Across the three GEA methods used, there were differences in the number and identity of SNPs associated with each climate variable. SNP associations for both species were found to be distributed throughout the genome, across each of the 11 chromosomes, with SNPs categorised into either gene space, regulatory (up to 1 Kb upstream of a gene), or non-genic regions. Across the three GEA methods, each GEA identified different suites of SNPs associated with the climate variables. However, we observed much congruence across any two of the methods utilised. Only one SNP for *E. blakelyi* (X06238 for annual aridity index, AAI) and four for *E. tereticornis* (X02631, X03997, and X04842 for temperature seasonality, T_SN_, and X11916 for maximum temperature of the warmest month, T_Max_) was identified by all GEA methods.

Of the 328 SNPs identified for *E. blakelyi*, from any one of the GEA methods, we observed the most associations for P_WARMQ_ (BIO18) (n=91), and 75 SNPs detected for both T_MA_ (BIO1) and AAI, respectively. Meanwhile, for *E. tereticornis*, we observed 130 SNPs for T_SN_ (BIO4), 99 SNPs for P_WARMQ_ (BIO18), and 87 SNPs for T_Min_ (BIO6).

For both species, we mapped these SNPs across the genome and linked them to the functional annotation. The proportion of genic SNPs identified from the GEA methods were 18.90% for *E. blakelyi* and 12.69% for *E. tereticornis*. For both species, however, we found that the most SNPs within genes to be for T_MA_ (28 vs 29 SNPs of the 75 SNPs identified via the GEAs accounting for 37.33% and 38.67% respectively; Table S5).

When we look at the number of shared SNPs among both species, we observe just 16 SNPs (2.19%) (Figure 2), with just two of these 16 SNPs associated with the same climate variate (P_DM_; BIO14) with a further two additional SNPs identified on the same sequencing fragment, but the SNP differs between species.

**Figure 2:**
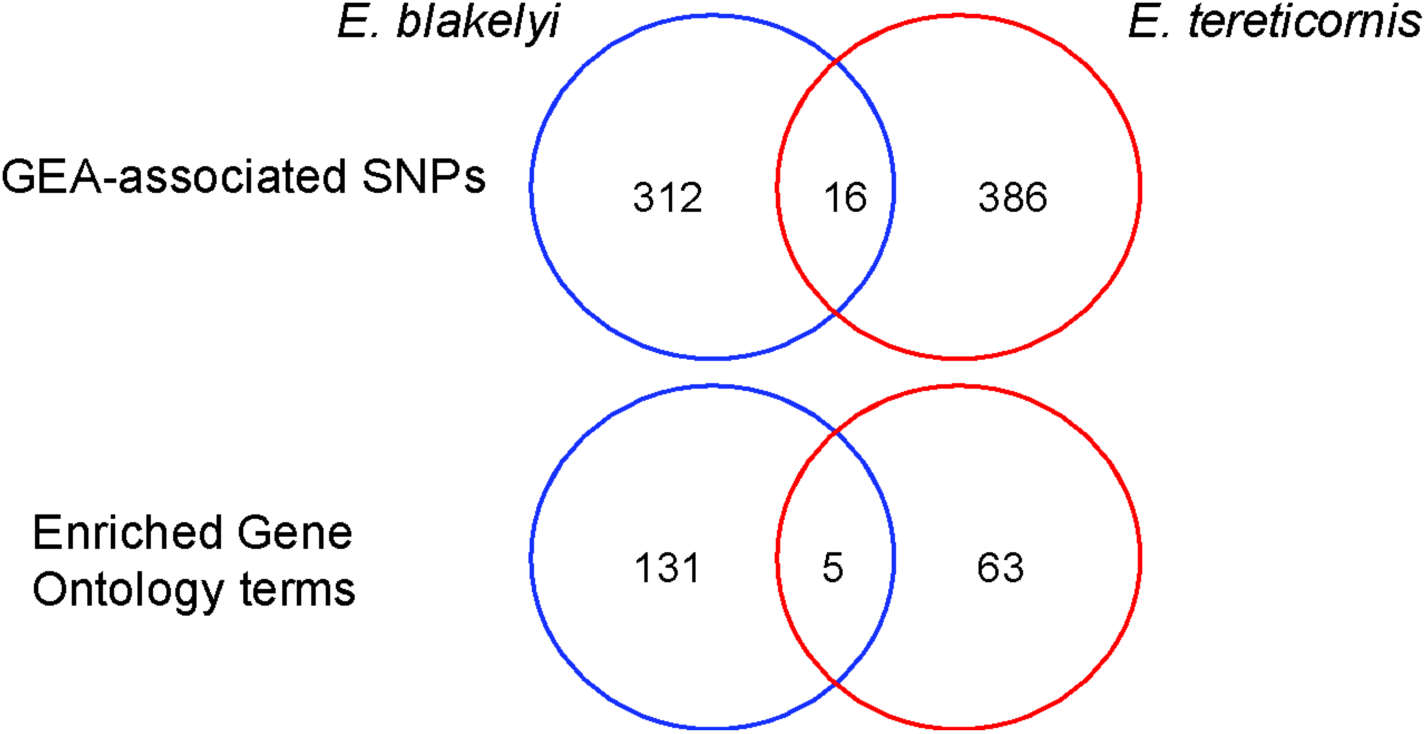
Venn diagrams showing total number of unique SNPs identified within each species, the intersection of shared SNPs identified within both species and the number of independently identified gene ontology terms per species and those shared. Blue are *E. blakelyi*; red are *E. tereticornis*. Enriched GO terms include cellular components, molecular function, and biological process.

### Generalised Dissimilarity Modelling

The SNPs for each climate variable were analysed in GDMs to determine their allelic turnover across climatic space (Figure 3). For two variables (T_Min_ and P_WARMQ_) we found that *E. tereticornis* had no observable allelic turnover over the climate space. All SNPs possessed a variable turnover (deviance explained) across the eight climate variables tested, with only five SNPs exhibiting a turnover greater than 0.15. Of these five SNPs, four belonged to *E. blakelyi* and one to *E. tereticornis*. For *E. blakelyi*, we found one SNP among each of the following climate variables: AAI (SNP X11717 identified as non-genic, allelic turnover of 0.25), P_MA_ (SNP X06793 identified as genic and a turnover of 0.15), P_WARMQ_ (SNP X00091 identified as genic, turnover of 0.19), and T_SN_ (SNP X08267, genic and a 0.19 allelic turnover). While SNP X04213 (regulatory) for *E. tereticornis* experienced a shift of 0.21 across the P_DM_ gradient. The majority of the SNPs possessed an allelic turnover of less than 0.1.

**Figure 3:**
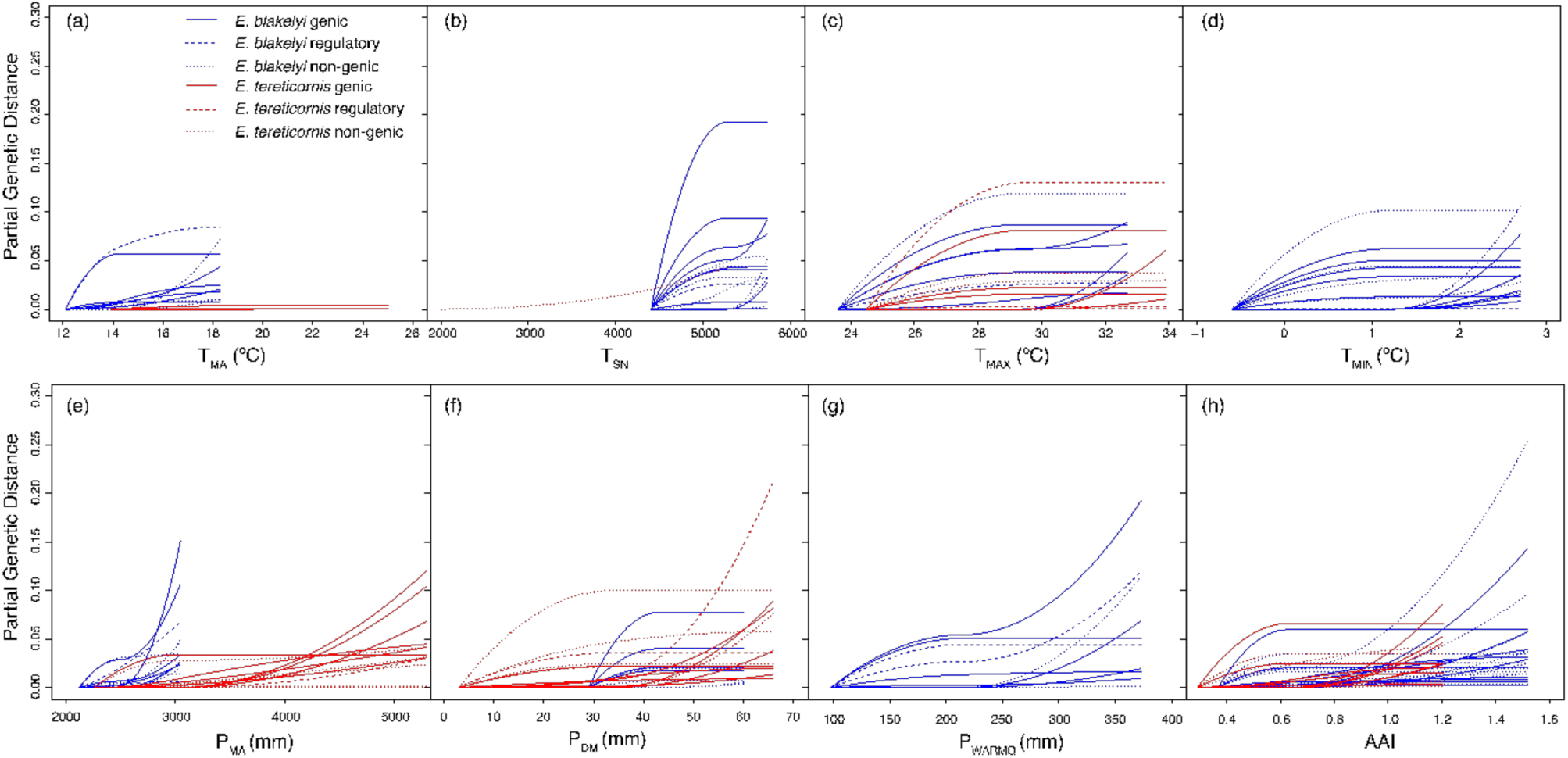
Spline plots of allelic turnover of genic, regulatory, and non-genic SNPs for *E. blakelyi* (blue) and *E. tereticornis* (red) for (a) T_MA_, (b) T_SN_, (c) T_Max_, (d) T_Min_, (e) P_MA_, (f) P_DM_, (g) P_WARMQ_, and (h) AAI. Solid lines indicate SNPs within genes, dashed lines for regulatory SNPs, and dotted lines for non-genic SNPs.

We also ran the GDM for groups of SNPs in the three enriched GO terms (see Table 1 for summary) common to both species. Two of the enriched GO terms (oxidoreductase activity, GO:0016491, and tetrapyrrole binding, GO:0046906) fell under molecular function (MF), while the third, oxidation-reduction process (GO:0055114) was under biological process (BP). It is important to note that across for the oxidoreductase activity and oxidation-reduction process enriched processes, for *E. blakelyi* there are 14 SNPs in common, and for *E. tereticornis* there are 28 SNPs in common, which results in a near identical distribution, thus we only show the maps for the biological process oxidation-reduction process. For *E. blakelyi*, we observe an east-west shift of the allele frequencies for both oxidoreductase activity and oxidation-reduction process using 15 and 17 SNPs respectively, while for *E. tereticornis*, (using 28 and 34 SNPs, respectively) we note a shift in its southern distribution and a less abrupt east-west shift in the middle of its distribution (Figure 4a-b). For the third GDM map (Figure 4c-d), tetrapyrrole binding, we observe a north-south shift from five SNPs for *E. blakelyi* and a latitudinal shift between NSW and QLD with further isolated regions along the coast and far north of its distribution for *E. tereticornis* compared to its other GDMs from 8 SNPs. For *E. tereticornis*, we observe areas of dissimilarity largely in the southern areas of its distribution, and isolated zones up the central coast, with an additional zone in the far north of its distribution. Within the oxidation-reduction process GO term we identify two *E. grandis* CDS fragments in common across both species; Eucgr.F03957 for the gene NDH-dependent cyclic electron flow 1, where the sequences overlap within the gene, and coproporphyrinogen III oxidase, where the sequences for each species largely do not overlap.

**Table 1:** Number of genic or regulatory SNPs for the enriched gene ontology terms of oxidation-reduction process and tetrapyrrole binding for *E. blakelyi* and *E. tereticornis*. Regulatory SNPs are defined as those ≤1 Kb upstream of a gene.

| Species | Oxidation-Reduction process |  | Tetrapyrrole binding |  |
| --- | --- | --- | --- | --- |
|  | Genic | Regulatory | Genic | Regulatory |
| *E. blakelyi* | 17 | 0 | 5 | 0 |
| *E. tereticornis* | 32 | 2 | 8 | 0 |

**Figure 4:**
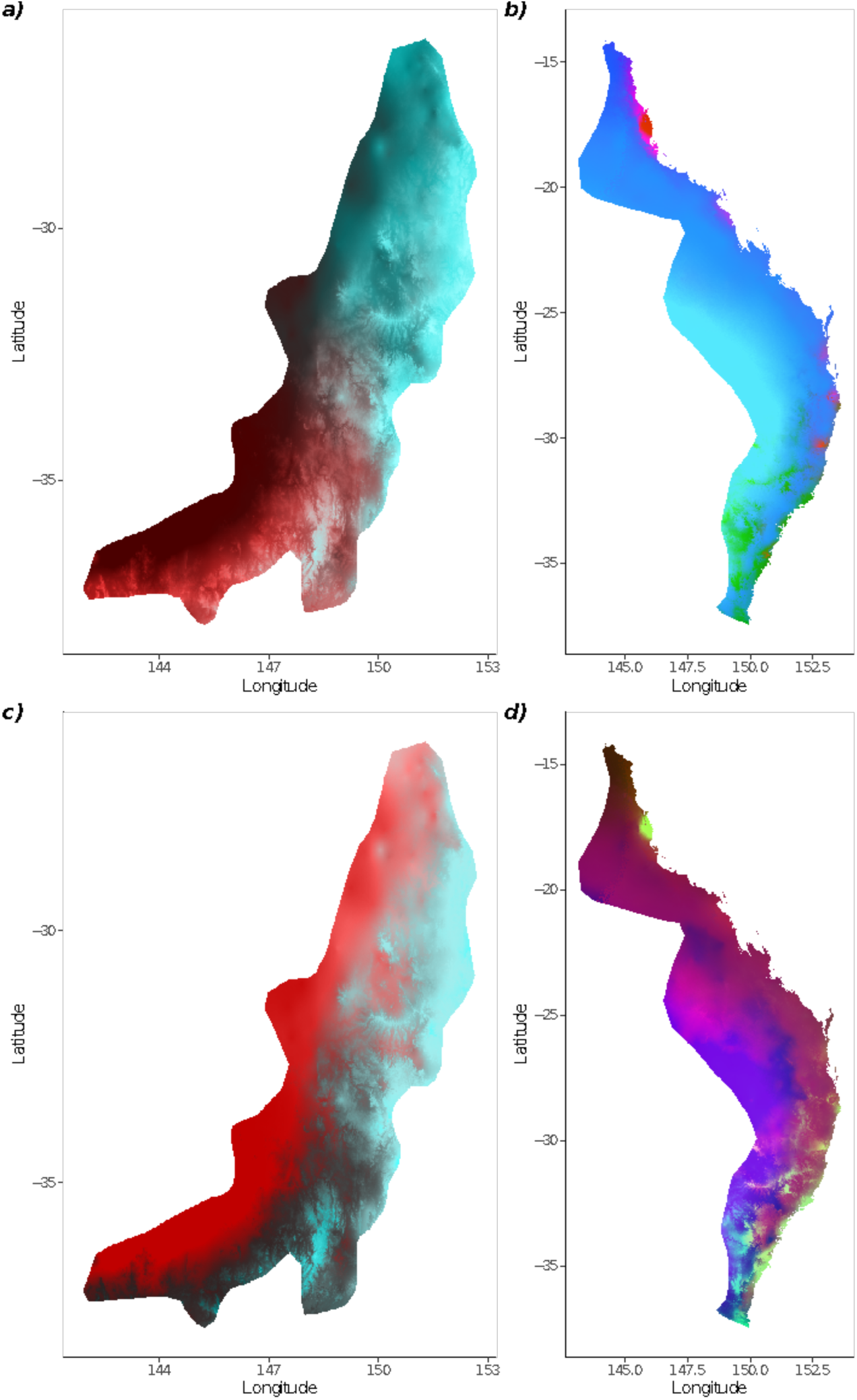
General dissimilarity modelling of *E. blakelyi* (a), c)) and *E. tereticornis* (b), d)) throughout their natural distributions. Top panels (a and b) show the allelic shifts for oxidation-reduction process, and bottom panels (c and d) show the changes for tetrapyrrole binding. Different scales used for each species’ maps since distributions of each species are not equal, and are on opposite sides of the Great Dividing Range.

Under future predictions with a 2070 climate outcome (Figure 5), for *E. tereticornis*, we identify populations along the coast (Figure 5b) and in New South Wales (NSW) (Figure 5d) that are projected to require the greatest amount of genomic turnover to keep up with a changing climate, in the worst-case scenario across both gene ontologies. Within the NSW region, the region most apparent at risk for the oxidation-reduction process GO term, also encompasses the critically endangered Cumberland Plain woodland ecological community within the western Sydney region in south-eastern Australia. For *E. blakelyi*, under future projections of both GO terms (Figure 5a, 5c), the furthest north, and the far southeast to be most at risk of population decline.

**Figure 5:**
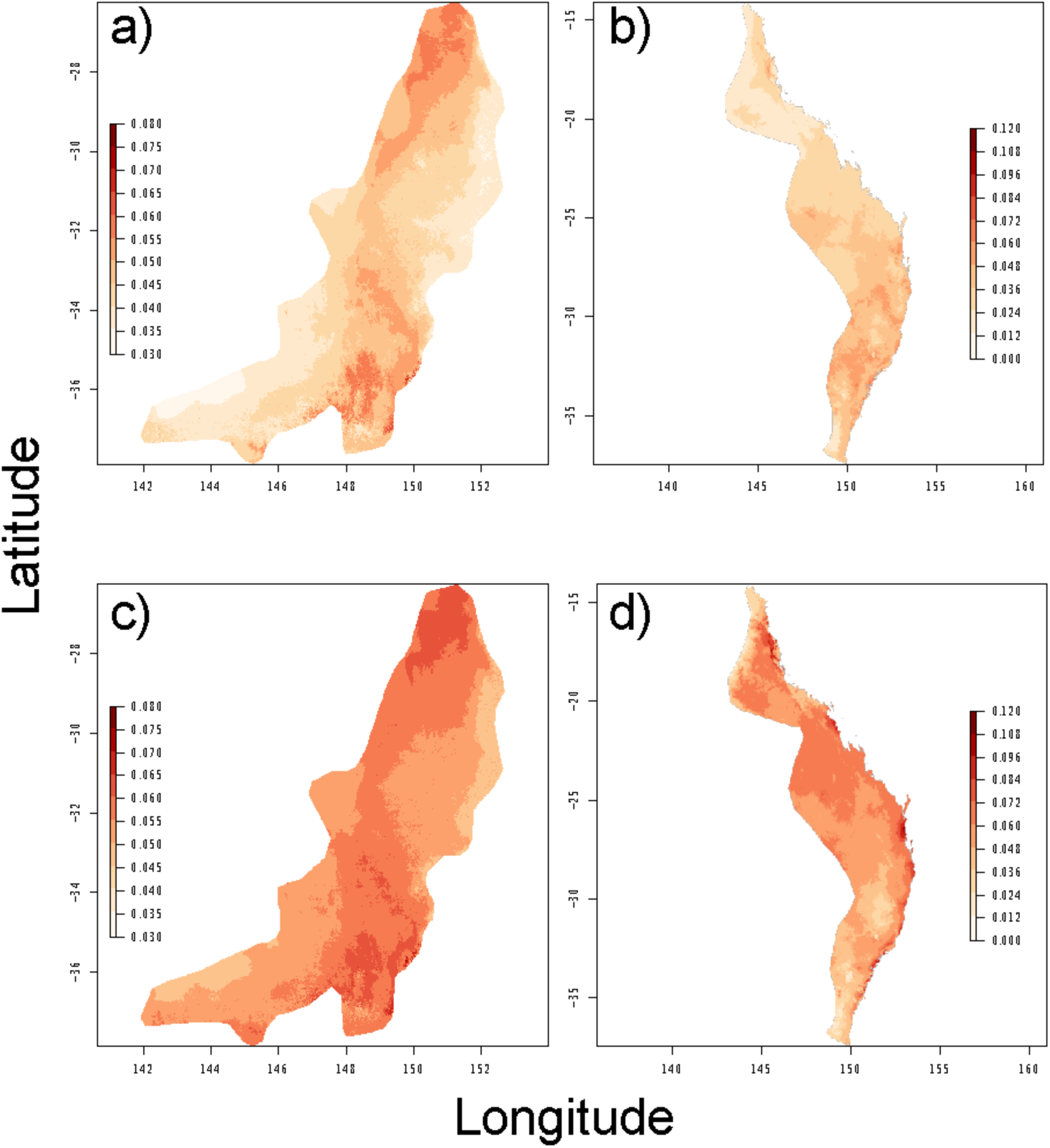
The predicted genomic vulnerability for *Eucalyptus blakelyi* (a, c), and *E. tereticornis* (b, d) based on comparing the GDM model projected on current climatic variables with the GDM model projected onto the predicted environmental conditions for 2070. Panels a) and b) are based on the SNPs associated with the oxidation-reduction process GO term, and panels c) and d) for the tetrapyrrole binding GO term. The greater the difference (dark red) between the two models, the greater amount of genomic change is required to keep up with predicted climate and environmental conditions.

## DISCUSSION

Our study was designed and set out to provide a comparison of the genomic adaptation to climate among two widespread red gum *Eucalyptus* species and estimate their capacity to respond under projected climates. We provide support for our first hypothesis with strong genetic signatures of local adaptation to climate-of-origin in both species at three hierarchical genomic levels (SNPs, genes, and gene ontology functions). In contrast to our hypothesis (2) that the two species would differ in the climate predictors explaining the most variation, we found that temperature was an important predictor of adaptation in both species. While we support hypothesis 3 with evidence for gene functions underlying adaptation, populations at the species climate margins were not a greater risk under future scenarios rejecting our last hypothesis. Understanding how these foundational species are putatively adapted to their climate and which populations are most vulnerable to climate change is critical for the maintenance and success of these trees in the future.

Eucalypts, like many other widespread tree species, exhibit low population genetic differentiation throughout their distributions, often with an *F*_ST_ of <0.08, as a result of high levels of gene flow, a preference for outcrossing, and connectivity between populations (Eldridge et al., 1993; Potts & Wiltshere, 1997). Our data on these red gum eucalypts show relatively low genetic differentiation in *E. blakelyi* 0.089 and *E. tereticornis* 0.129; the level of population structure is higher than non-red gum eucalypt species, but lower than has previously been observed in the river red gum *E. camaldulensis* (*F*_ST_ 0.17; Dillon et al., 2014). Similarly, a previous study of *E. tereticornis* found a greater-than-average F_ST_ of 0.099 using microsatellites, albeit with fewer populations studied (n=9) than within this study (Rymer unpubl.).

Previous studies investigating genomic adaptation to climate in eucalypt species have identified a greater proportion of SNPs associated with temperature variables than precipitation or aridity (Ahrens et al., 2019; Dillon et al., 2014; Jordan et al., 2017; Steane et al., 2014, 2017), as we have for *E. tereticornis*, but not for *E. blakelyi* where a greater proportion was found for rainfall-driven SNPs. Further, enriched gene ontology functions were precipitation-driven (BIO18 and AAI) for *E. blakelyi*, while temperature and precipitation variables were significant for *E. tereticornis*. Thus, we partially accept our hypothesis that temperature was the main driver of adaptation for *E. tereticornis* and accept our hypothesis that precipitation variables are more important for *E. blakelyi*. These differences in signatures of selection between the two species are likely due to the considerable differences in their precipitation variables (Figure S1), creating different adaptive requirements for their respective environments. This pattern is indicative of patterns of local adaptation among species with the accumulation of temperature- or precipitation-related SNPs. However, the temperature-related SNPs may partially be due to false positives of unaccounted for latitudinal structure within each species, despite our attempts at controlling for population structure through spatial autocorrelation and effective sample size (See supplementary information and Table S3).

Local adaptation to climate may arise through natural selection acting upon a number of small-effect loci that may facilitate the emergence population structure within the system (Orr, 2005). As local climates begin to change, these specific adaptations held by local populations may become problematic, subsequently decreasing species fitness. By 2100, mean global temperature increases are projected to exceed 1.5°C (IPCC, 2013), with heatwaves and extreme precipitation events e.g. flooding and droughts, projected to increase in their frequency and duration, including across eastern Australia (CSIRO (Australia) & Bureau of Meteorology, 2015; IPCC, 2014). These shifts however, are not anticipated to be uniform, with the northern tropics projected to receive an increase in annual, but variable rainfall, while the southeast may expect a reduction (Sillmann et al., 2013; Teng et al., 2012). As rainfall patterns and temperatures proceed to change in eastern Australia, this may cause populations of both species to become maladapted in their current location. Recent studies have observed physiological and hydraulic adaptation to climate and particularly along aridity gradients (Drake et al., 2015; Li et al., 2018; McLean et al., 2014; Steane et al., 2015; 2017), and also the phenotypic plasticity of physiological processes under varying temperature in eucalypts (Drake et al., 2017). Here, we identified a number of putatively adaptive SNPs across eight climate variables for both eucalypts, mostly within genic (coding) regions, and some within regulatory regions. SNPs within these genomic regions can play a role in phenotypic divergence that may lead to adaptation, either through the direct changing of a phenotype, or via a change in expression in the response of stimuli.

Our data supports the independent evolution and adaptation to climate in these two closely-related species in that we observe few shared SNPs and few common GO terms between them, thus supporting our hypothesis that the SNPs will differ between the species. Though hybridisation among eucalypts, and between these two species (Brooker & Slee, 2000; House, 1997; Klaphake, 2012), may inhibit genomic differentiation, our data provides evidence for independent evolution in these closely related species. The different climate envelopes these species exhibit appear to have driven adaptation to distinct variables (Figure 3). We find that climate explains up to 25% of the allelic turnover in these species. However, many SNPs show low levels (<15% partial genetic distance) of turnover with specific climate variables and SNPs shift in frequency at different stages along the climate envelope. Adaptation to climate here, like other tree species, is likely driven by many loci of small effect (Savolainen et al., 2013) rather than few loci of large effect. Therefore, the characterisation of these suite of SNPs and their associated genes allows for further analysis and investigation of the influence they may have upon climate adaptation. Previous studies have performed GDM analyses on SNP datasets to categorise the allele frequency shift (or allelic turnover), and genomic vulnerability across the landscape (Ahrens et al., 2019; Cao et al., 2020; Fitzpatrick & Keller, 2015; Supple et al., 2018). Our data did not highlight one or few SNPs of large allelic turnover across the landscape, therefore we subsequently performed this analysis on a collection of SNPs associated with the common enriched GO term found among both eucalypts.

### Annotation, gene function, and genomic vulnerability of enriched GO terms

We found significant SNP-climate associations within gene space, regulatory regions, and non-genic space. Both the molecular function of oxidoreductase activity and the biological process of oxidation-reduction process possess a large number of common SNPs between them, and as such are associated with the same or similar function. The process itself involves the enzymatic catalysis of electron transfer from one compound to another, often associated with photosynthesis and respiration and utilising an NAD+ or NADH cofactor. Here we find two SNPs within genes associated with photosynthesis in *E. blakelyi* (SNP X01686 associated with P_WARMQ_, and SNP X12218 associated with T_SN_) and one SNP (X05388 associated with T_MA_) in a gene that may be associated with cold acclimation (Dyson et al., 2016). We also note one SNP (X01686 associated with P_WARMQ_) within the cytokinin oxidase 3 gene; involved in the oxidation of cytokinins (Werner et al., 2003) in *Arabidopsis*. Cytokinins are a group of phytohormones involved in growth and development (Zubo et al., 2017) though may influence or impact other phytohormones involved in both abiotic and biotic stress responses (Großkinsky et al., 2016). For *E. tereticornis*, we detect two SNPs (X01577 associated with several climate variables - T_SN_, T_Min_, P_MA_, P_WARMQ_, and AAI, and SNP X07527 associated with both T_Max_ and T_Min_) associated with genes within the jasmonic acid pathway, a key plant hormone in herbivory defence and environmental responses (Creelman & Mullet, 1995; Du et al., 2013; J. Li et al., 2018). In addition to these SNPs and genes or proteins specific to these species within this GO term, we detected two proteins found in both species. The first, NDH-dependent cyclic electron flow 1 (SNP X02482 for *E. blakelyi* with AAI, and SNP X00150 for *E. tereticornis* associated with P_WARMQ_), which consist of overlapping sequences from both species. This gene is involved in ATP synthesis through the Photosystem I phase of photosynthesis (Takabayashi et al., 2009). The second protein, coproporphyrinogen III oxidase (with SNP X10124 for *E. blakelyi* associated with P_MA_ and AAI, and for *E. tereticornis*, SNP X11913 with the drier climate variables of T_Max_ and P_DM_) whose sequences for both species, largely do not overlap within this gene space. These SNPs and the subsequent protein, despite being a key (intermediate) enzyme in the biosynthesis of chlorophyll and hemes (Ishikawa et al., 2001; Santana et al., 2002), within the tetrapyrrole pathway, were not detected among the tetrapyrrole binding GO term. The molecular function of oxidoreductase activity has been found to be independently upregulated in *E. globulus* and *E. urograndis* seedlings when grown under varying temperature regimes for 30 days (Araújo et al., 2018) and both up- and down-regulated in *E. nitens* when under cold acclimated and de-acclimated conditions (Gaete-Loyola et al., 2017). Since this molecular function was observed to be up, under these conditions for *E. nitens*, this may indicate that several of the genes associated in this process (electron transfer) may have a form of “adaptive molecular memory” that may well be epigenetic. Further investigation into these genes among *E. blakelyi* and *E. tereticornis* are required to determine whether they are important in these species.

Biosynthesis of tetrapyrroles are essential for living organisms as they form the basis of hemes, key for oxidative and energy functions, and of chlorophyll, fundamental in photosynthetic activity (Cihlář et al., 2016; Kořený et al., 2011). In fact, it has been shown that these tetrapyrroles are essential for photosynthetic pathways (Papenbrock et al., 1999). Under this enriched GO term, despite identifying a handful of associated SNPs we were able to detect relevant genes. For *E. blakelyi* we identify cytochrome P450 (associated with SNP X01457) and light harvesting complex of photosystem II 5 (associated with SNP X10903) genes for P_WARMQ_ and AAI respectively. Cytochrome P450s are known to be involved with a number of plant biosynthetic pathways including those for plant defence, hormones, and secondary metabolites (Mizutani & Sato, 2011; Schuler & Werck-Reichhart, 2003), while the latter gene is essential in the light-dependent reaction in photosynthesis (Loll et al., 2005). This same gene (light harvesting complex of photosystem II 5) has also been found to be differentially expressed among three provenances of *E. camaldulensis* seedlings (humid tropics, dry tropics, and semi-arid) following placement under a water-stress treatment (Thumma et al., 2012). It appears that this gene is fundamental in the adaptation of red gum eucalypts, with the greatest provision for this adaptation within *E. blakelyi* are among populations with an even distribution of the X10903 allele frequencies. For *E. tereticornis*, we identified two SNPs, but three CDS regions associated with a peroxidase superfamily protein (SNP X05871 associated with P_WARMQ_, and SNP X06496 associated with T_MA_). This gene is involved in a number of metabolic pathways that covers removal of hydrogen peroxide, biosynthesis and degradation of lignin, environmental stress response e.g. oxidative stress and pathogen attack, and the oxidation of toxic reductants (Valério et al., 2004; Welinder et al., 2002). The latter, through the accumulation of reactive oxygen species (ROS), particularly under environmental stressors, are highly toxic that result in membrane and chloroplast degradation, affecting metabolic pathways, restricting and leading to a cessation of cellular function and growth (Cakmak, 2005; Dat et al., 2000; Reddy et al., 2004). Further to the common GO terms detected within both species, we identified a number of enriched terms that are species-specific. A subsetted list of these, the associated SNPs and climate variables, and proteins may also be viewed in the supplementary information.

### Vulnerability to climate change, and conservation implications

Predicting the populations and species that may be most at risk of future declines is fundamental in effective conservation strategies. Climate change is shifting the suitability of local populations to the environment they are exposed to now and with greater mismatches into the future (Browne et al., 2019). Projections of the extent of the genomic changes required to maintain the genotype-environment relationships under future climate scenarios, and is termed genomic vulnerability (Bay et al., 2018; Fitzpatrick & Keller, 2015; Rellstab et al., 2015; Rhoné et al., 2020; Ruegg et al., 2018; Waldvogel et al., 2020). Through the use of the GDM model, the genomic vulnerability of these SNPs among the enriched photosynthetic-process gene ontology terms described above, across the landscape can be characterised. For *Eucalyptus blakelyi*, we found the northernmost and south-eastern populations to be most vulnerable in projections of the SNPs under both GO terms. Both regions experience relatively low rainfall during the driest month but have an inverse relationship to one another in aridity; with low aridity in the north, and higher aridity in the mesic southeast. Neither region represents the climatic extremes for this species, but are the distribution extremes and perhaps the most isolated. For *E. blakelyi*, these regions of vulnerability identified also resemble the lower elevations of its distribution. Similarly for *E. tereticornis*, the regions exhibiting the greatest genomic vulnerability under a future climate scenario are not among the current climatic extremes, therefore we may reject our final hypothesis for both species. This approach has also been utilised to link that early and late flowering varieties of pearl millet in West Africa are genomically vulnerable to decline (Rhoné et al., 2020), and that populations of the yellow warbler in North America that have already experienced declines also require the greatest allele frequency shifts to prevent further declines due to climate change Bay et al., 2018). Based on these projections, it would appear the southernmost distribution in NSW for *Eucalyptus tereticornis* to be least affected under both GO term scenarios, and may be the most appropriate seed-source location. For *E. blakelyi* however, regions projected to be least at risk are those at the edge of its distribution in central Victoria and the New England Tablelands. For the protection and maintenance of these species, additional studies on their physiological responses to environmental change must be undertaken. However, understanding the adaptive genomics of these species, and their genomic diversity is the first step in this process.

Changes in land-use, through agriculture and urbanisation, may further impact on the ability for populations to keep pace with climate change. A reduction in population density and connectivity can limit adaptive capacity associated with standing genomic variation and gene flow. The Sydney Basin and Hunter regions may represent the most isolated or fragmented populations. The critically endangered Cumberland Plain Woodland is also located within the Sydney Basin, and is predicted to require approximately 6% genomic turnover to keep track with future climate shifts. Given continued urban growth pressures with biodiversity offsetting in this region mandated to conserve and restore Cumberland Plain Woodland it will be important to actively manage red gum populations (CPCP, NSW Department of Planning, Industry and Environment (2020)). Adaptive management strategies, including assisted gen migration and establishment of diverse translocated populations, may be informed by genomic predictions of climate adaptation now and into the future.

## CONCLUSION

The two red gum species exhibit strong patterns of adaptation to their local climates. However, empirical studies will be required to validate whether these adaptations translate into trait or physiological variation among provenances. Additionally, the suite of candidate SNPs associated with climate adaptation appear to have occurred through divergent adaptation despite the close-relatedness of these species. Based on our generalised dissimilarity modelling across the climatic landscape, we found that precipitation during the warmest quarter and annual aridity index to be most important for *E. blakelyi*. Meanwhile for *E. tereticornis*, these climate variables plus mean annual temperature, temperature seasonality, mean annual precipitation, and precipitation of the driest month were key for driving adaptive genomic variation. The candidate SNPs are found in key photosynthetic and plant hormone pathways, which may buffer against environmental stressors. Furthermore, we identify genomic vulnerability for both species in marginal populations that may require active management in the future. To mitigate population declines of these foundation species, strategies such as assisted gene migration may be employed to supplement populations and enhance the frequency of adaptive alleles.

