## Supplementary File 1 for "Closely Related Tree Species with Overlapping Ranges Exhibit Divergent Adaptation to Climate"

### SUPPLEMENTARY INFORMATION

#### Results

*Environmental Associations*

We tested for spatial autocorrelation (Table S3) among the climate variables using Moran's *I* to test the power of the sampling design, and their effective sample sizes (ESS) in the presence of SA. For *E. blakelyi*, we detected low climate spatial autocorrelation across all populations for seven of the eight variables used in this study, with ESS remaining large across climate variables for this species (Table S3). For *E. tereticornis*, we detected strong climate spatial autocorrelation across all populations for each climate variable, thus observing lower ESS values than for *E. blakelyi*. The greatest ESS for *E. blakelyi* were observed in the $P_{MA}$, $P_{DM}$ and $T_{Min}$, while AAI, $T_{Max}$, and $P_{DM}$ represented the highest ESS (and subsequently lowest spatial autocorrelation). This inverse relationship between spatial autocorrelation and ESS occurs through the accumulation of increasing redundant information in the climate variables between locations. Simply, as spatial autocorrelation increases, ESS decreases. As such, the narrower longitudinal climate envelope in the distribution for *E. tereticornis* between the Great Dividing Range (GDR) and the coast constrains its ESS compared to *E. blakelyi*. Pearson's correlation coefficients (Table S4) of the eight climate variables showed differences in pairwise *r* values for both species, with strong relationships observed for one species were low in the other e.g. BIO1 vs BIO18, *r* = 0.728 for *E. tereticornis* and *r* = 0.051 for *E. blakelyi*, and similarly BIO4 vs BIO5 *r* = 0.852 for *E. blakelyi* and *r* = 0.009 for *E. tereticornis* (Table S4).

Table S1: Populations, latitudes, longitudes, and climatic variables used in this study for *Eucalyptus tereticornis*; $T_{MA}$ (mean annual temperature), $T_{SN}$ (mean temperature seasonality), $T_{Max}$ (mean maximum temperature of the warmest month), $T_{Min}$ (mean minimum temperature of the coldest month), $P_{MA}$ (mean annual precipitation), $P_{DM}$ (mean precipitation of the driest month), $P_{WARMQ}$ (mean precipitation of the warmest quarter), AAI (mean annual aridity index).

| Population | Longitude | Latitude | $T_{MA}$ | $T_{SN}$ | $T_{Max}$ | $T_{Min}$ | $P_{MA}$ | $P_{DM}$ | $P_{WARMQ}$ | AAI |
|---|---|---|---|---|---|---|---|---|---|---|
| ACCR | 152.27646 | -28.38187 | 16.3 | 4416 | 28.1 | 2.8 | 910 | 38 | 345 | 0.634 |
| ANAN | 145.17188 | -15.67965 | 25 | 2009 | 31.7 | 17.6 | 1814 | 20 | 933 | 0.945 |
| APPI | 150.78339 | -34.19791 | 16.1 | 4034 | 26.7 | 3.9 | 1072 | 47 | 322 | 0.563 |
| BOGN | 149.72536 | -26.46732 | 19.6 | 5315 | 33.1 | 3.9 | 631 | 27 | 251 | 0.339 |
| BROK | 151.09378 | -32.72147 | 17.3 | 4577 | 29.8 | 3.9 | 685 | 28 | 246 | 0.571 |
| BRSF | 151.17683 | -28.20235 | 18.6 | 5144 | 32.1 | 3.5 | 660 | 30 | 244 | 0.339 |
| CHMB | 152.72098 | -29.80756 | 18.4 | 4066 | 29.1 | 5.3 | 1297 | 39 | 521 | 0.628 |
| CRNP | 150.15155 | -35.6696 | 15.9 | 3545 | 25 | 5.7 | 1068 | 59 | 321 | 0.801 |
| DACR | 146.04323 | -18.55243 | 23.4 | 2982 | 31.4 | 12.8 | 1819 | 27 | 928 | 1.027 |
| DESF | 152.61844 | -25.61844 | 21.3 | 3489 | 30.3 | 9.9 | 1104 | 31 | 458 | 0.642 |
| FOMI | 144.85501 | -18.08131 | 21.4 | 3185 | 31.6 | 9.2 | 693 | 3 | 329 | 0.434 |
| GATN | 152.24156 | -27.54052 | 19 | 4396 | 30.7 | 4.8 | 874 | 32 | 352 | 0.451 |
| GIHI | 153.27673 | -28.81917 | 19.4 | 3695 | 28.9 | 7.4 | 1427 | 42 | 506 | 0.941 |
| GLEN | 153.06413 | -29.84003 | 18.8 | 3741 | 28 | 6.8 | 1390 | 44 | 482 | 0.803 |
| GRMO | 149.03771 | -21.17854 | 22.4 | 3436 | 30.9 | 10.9 | 1639 | 17 | 869 | 0.777 |
| HAST | 145.47812 | -17.29574 | 20.6 | 2728 | 29.2 | 10.7 | 1706 | 34 | 803 | 0.903 |
| HUSF | 148.85493 | -24.16634 | 21.3 | 4711 | 33.9 | 6.1 | 680 | 21 | 319 | 0.297 |
| JELL | 150.38901 | -34.36928 | 13.9 | 4350 | 25.9 | 1.7 | 942 | 46 | 276 | 0.637 |
| KUKU | 151.45599 | -32.82007 | 17.7 | 4210 | 28.7 | 5.3 | 879 | 36 | 295 | 0.597 |
| LEJA | 152.15108 | -29.71363 | 15.7 | 4458 | 27.6 | 2.4 | 1042 | 41 | 439 | 0.543 |
| MIDL | 148.66895 | -23.07637 | 22.1 | 4278 | 33.5 | 7.9 | 632 | 16 | 303 | 0.292 |
| MOCR | 149.64235 | -36.66888 | 14.7 | 3941 | 25.6 | 3.4 | 887 | 49 | 268 | 0.632 |
| MOHO | 148.24188 | -25.33656 | 19.1 | 5322 | 33 | 2.8 | 669 | 18 | 292 | 0.333 |
| MUHY | 144.72665 | -16.35493 | 23.4 | 2329 | 32.1 | 13.8 | 1286 | 10 | 539 | 0.521 |
| MUNG | 145.25072 | -15.70315 | 24.2 | 2060 | 31 | 16.9 | 1846 | 23 | 935 | 0.989 |
| NORO | 151.35593 | -32.77314 | 17.6 | 4346 | 29.3 | 4.8 | 798 | 31 | 278 | 0.537 |
| NWRA | 150.61243 | -34.91884 | 16.6 | 3442 | 25.4 | 7.1 | 1267 | 66 | 397 | 0.779 |
| NYMB | 152.73491 | -29.98572 | 17.9 | 4066 | 28.5 | 5 | 1463 | 44 | 581 | 0.726 |
| POCU | 150.53134 | -23.43509 | 22.3 | 3786 | 31.6 | 9.7 | 869 | 21 | 415 | 0.43 |
| RICH | 150.73746 | -33.61771 | 17.2 | 4501 | 29 | 3.3 | 894 | 39 | 309 | 0.595 |
| ROSS | 145.22842 | -15.75363 | 24.5 | 2055 | 31.3 | 17 | 1827 | 21 | 932 | 0.989 |
| RVTE | 152.27103 | -28.63852 | 17.8 | 4381 | 29.7 | 4.2 | 880 | 34 | 352 | 0.513 |
| SCON | 150.79063 | -32.01361 | 16.6 | 4972 | 30.4 | 2.6 | 731 | 39 | 266 | 0.434 |
| SMRP | 145.67912 | -16.79846 | 24.4 | 2341 | 31.4 | 16.2 | 2146 | 33 | 1067 | 1.006 |
| TATH | 149.98056 | -36.73713 | 15.2 | 3511 | 24.5 | 5.1 | 922 | 49 | 273 | 0.705 |
| TOMA | 151.55084 | -31.78715 | 14.2 | 4545 | 26.4 | 1.4 | 958 | 47 | 345 | 0.722 |
| TUSF | 145.46860 | -17.53347 | 19.2 | 2852 | 28.3 | 8.8 | 1364 | 29 | 647 | 1.203 |
| WEBU | 153.08736 | -29.08736 | 19.3 | 3768 | 29 | 7.1 | 1253 | 38 | 456 | 0.699 |

Table S2: Populations, latitudes, longitudes, and climatic variables used in this study for *Eucalyptus blakelyi*; $T_{MA}$ (mean annual temperature), $T_{SN}$ (mean temperature seasonality), $T_{Max}$ (mean maximum temperature of the warmest month), $T_{Min}$ (mean minimum temperature of the coldest month), $P_{MA}$ (mean annual precipitation), $P_{DM}$ (mean precipitation of the driest month), $P_{WARMQ}$ (mean precipitation of the warmest quarter), AAI (mean annual aridity index).

| Population | Longitude | Latitude | $T_{MA}$ | $T_{SN}$ | $T_{Max}$ | $T_{Min}$ | $P_{MA}$ | $P_{DM}$ | $P_{WARMQ}$ | AAI |
|---|---|---|---|---|---|---|---|---|---|---|
| ASCA | 152.00598 | -30.35624 | 12.1 | 4402 | 23.6 | 0 | 912 | 44 | 356 | 0.677 |
| BEBU | 150.02957 | -33.25019 | 12.2 | 4784 | 25.5 | 0.3 | 805 | 51 | 252 | 0.673 |
| BEHP | 146.68135 | -36.34818 | 12.4 | 5206 | 27.4 | 0.8 | 1024 | 38 | 160 | 1.519 |
| BMRA | 147.20186 | -35.84277 | 14.4 | 5490 | 30.2 | 2.1 | 766 | 38 | 137 | 0.945 |
| BOOK | 148.63074 | -34.81331 | 13.8 | 5481 | 29.8 | 0.4 | 757 | 49 | 165 | 0.881 |
| CNBR | 149.1029 | -35.1649 | 12.4 | 5235 | 27.6 | -0.6 | 743 | 50 | 173 | 0.572 |
| CONP | 148.41368 | -33.73412 | 15.2 | 5578 | 31.3 | 1.8 | 722 | 50 | 179 | 0.607 |
| DDNP | 151.03464 | -28.88149 | 18.3 | 5368 | 32.7 | 2.7 | 693 | 33 | 262 | 0.368 |
| DNDE | 151.92319 | -29.59520 | 13.5 | 4517 | 25.4 | 0.7 | 956 | 49 | 373 | 0.683 |
| GLBN | 149.75439 | -34.74943 | 12.9 | 4793 | 27 | 0.7 | 701 | 44 | 189 | 0.559 |
| GNR | 149.28613 | -33.15732 | 13.9 | 5223 | 29 | 0.6 | 749 | 49 | 208 | 0.63 |
| GOOB | 148.38484 | -32.64029 | 16.1 | 5608 | 31.4 | 2.6 | 699 | 44 | 196 | 0.522 |
| KOYO | 143.68369 | -36.58689 | 14 | 4875 | 29.1 | 2.7 | 567 | 31 | 98 | 0.745 |
| LWDE | 145.44388 | -36.84907 | 13.3 | 4920 | 28.1 | 2.4 | 797 | 34 | 132 | 1.052 |
| MGNR | 149.83603 | -32.39433 | 14.8 | 5188 | 28.8 | 1.2 | 724 | 43 | 236 | 0.503 |
| MINP | 148.17950 | -35.24604 | 14.8 | 5484 | 31 | 1.3 | 836 | 43 | 162 | 0.948 |
| MOLR | 150.28082 | -30.34455 | 14.1 | 5542 | 28.8 | -0.5 | 1009 | 60 | 334 | 0.743 |
| MOTO | 151.25555 | -29.92684 | 14.2 | 5120 | 27.9 | -0.1 | 853 | 43 | 305 | 0.618 |
| MRR | 148.088 | -36.036 | 14.1 | 5355 | 30 | 1.2 | 891 | 44 | 171 | 1.1 |
| OSTR | 152.03815 | -28.42674 | 15.7 | 4574 | 27.7 | 2.1 | 814 | 36 | 310 | 0.54 |
| PLGA | 149.40080 | -30.93342 | 17 | 5654 | 32.3 | 2 | 723 | 43 | 252 | 0.415 |
| PREM | 149.89836 | -31.45145 | 16.9 | 5466 | 31.6 | 2.2 | 644 | 29 | 244 | 0.387 |
| PUD | 148.93430 | -34.58340 | 13 | 5328 | 28.7 | 0 | 747 | 52 | 172 | 0.712 |
| RORR | 151.47670 | -30.60239 | 13.2 | 4882 | 26.4 | -0.4 | 789 | 38 | 288 | 0.606 |
| STOC | 147.97269 | -34.55496 | 14.8 | 5729 | 31.2 | 1.6 | 695 | 42 | 154 | 0.698 |
| TEAM | 151.05838 | -31.54967 | 13.8 | 4925 | 27.3 | 0.4 | 868 | 50 | 284 | 0.672 |
| THUD | 148.22816 | -34.18013 | 15 | 5592 | 31.1 | 1.5 | 658 | 45 | 150 | 0.63 |
| ULAN | 149.79337 | -32.22064 | 15.7 | 5259 | 29.8 | 1.7 | 665 | 38 | 223 | 0.427 |
| UPDR | 150.59071 | -31.87082 | 14.3 | 5079 | 28 | 0.7 | 915 | 56 | 310 | 0.534 |
| WAGG | 147.37007 | -35.05303 | 15.7 | 5654 | 31.6 | 2.7 | 563 | 35 | 121 | 0.564 |
| WAOV | 146.20192 | -36.28669 | 14 | 5275 | 29.5 | 2.2 | 684 | 33 | 127 | 0.823 |
| WBNG | 150.75040 | -32.20449 | 16.5 | 4924 | 30.2 | 2.7 | 706 | 38 | 258 | 0.444 |
| WENR | 149.59604 | -31.71314 | 15.3 | 5444 | 30 | 1.1 | 719 | 46 | 237 | 0.462 |
| WSCA | 150.58452 | -29.65145 | 16.8 | 5575 | 31.8 | 0.7 | 712 | 34 | 255 | 0.416 |

Table S3: Spatial autocorrelation (Moran's *I*) and effective sample size (ESS) of the climate variables for *E. blakelyi* and *E. tereticornis*. BIO1 - mean annual temperature ($T_{MA}$), BIO4 - temperature seasonality ($T_{SN}$), BIO5 - mean maximum temperature of the warmest month ($T_{Max}$), BIO6 - mean minimum temperature of the coldest month ($T_{Min}$), BIO12 - mean annual precipitation ($P_{MA}$), BIO14 - (mean precipitation of the driest month ($P_{DM}$), BIO18 - mean precipitation of the warmest quarter ($P_{WARMQ}$), AAI (mean annual aridity index).

| | ***E. blakelyi*** | | | ***E. tereticornis*** | | |
|---|---|---|---|---|---|---|
| | Moran's *I* | p-value | ESS | Moran's *I* | p-value | ESS |
| **BIO1** | 0.06117644 | 0.01063081 | 31.0147598 | 0.59362 | 0.00 | 9.00186262 |
| **BIO4** | 0.1232449 | 1.95E-05 | 27.2668309 | 0.5955956 | 0.00 | 8.94552281 |
| **BIO5** | 0.05830512 | 0.01318787 | 31.1906963 | 0.3763398 | 3.48E-10 | 16.7013272 |
| **BIO6** | 0.04825381 | 0.03047144 | 31.8044365 | 0.6628551 | 0.00 | 7.15101278 |
| **BIO12** | 0.00553251 | 0.3150372 | 34.1217080 | 0.4129349 | 6.74E-12 | 15.1732798 |
| **BIO14** | 0.02461346 | 0.1266009 | 33.2023218 | 0.3934567 | 3.73E-11 | 15.9736385 |
| **BIO18** | 0.4066474 | 0.00 | 13.8686250 | 0.5397419 | 0.00 | 10.6243626 |
| **AAI** | 0.2170606 | 2.17E-13 | 22.1103348 | 0.2963448 | 4.64E-07 | 20.421982 |

Table S4: Pearson's correlation coefficient (*r*) of the eight bioclimatic variables

| | | **BIO1** | **BIO4** | **BIO5** | **BIO6** | **BIO12** | **BIO14** | **BIO18** | **AAI** |
|---|---|---|---|---|---|---|---|---|---|
| ***E. blakelyi*** | BIO1 | 1 | | | | | | | |
| | BIO4 | 0.4933032 | 1 | | | | | | |
| | BIO5 | 0.81610931 | 0.85207497 | 1 | | | | | |
| | BIO6 | 0.64641047 | 0.23346246 | 0.61049748 | 1 | | | | |
| | BIO12 | -0.4779868 | -0.3547137 | -0.5731604 | -0.5788728 | 1 | | | |
| | BIO14 | -0.4224953 | 0.00175864 | -0.323046 | -0.6611108 | 0.50934391 | 1 | | |
| | BIO18 | 0.05096782 | -0.4741371 | -0.4427923 | -0.4224934 | 0.53503578 | 0.32456516 | 1 | |
| | AAI | -0.572248 | -0.0436436 | -0.2479274 | 0.1222831 | 0.51038295 | 0.0254719 | -0.4029419 | 1 |
| ***E. tereticornis*** | BIO1 | 1 | | | | | | | |
| | BIO4 | -0.6320153 | 1 | | | | | | |
| | BIO5 | 0.75783099 | -0.0086481 | 1 | | | | | |
| | BIO6 | 0.89351342 | -0.8995645 | 0.40639553 | 1 | | | | |
| | BIO12 | 0.5550665 | -0.8240704 | -0.0079146 | 0.77731937 | 1 | | | |
| | BIO14 | -0.7394647 | 0.20800745 | -0.859932 | -0.4806222 | -0.0090745 | 1 | | |
| | BIO18 | 0.72803398 | -0.8117098 | 0.24457238 | 0.86123532 | 0.94933229 | -0.2747797 | 1 | |
| | AAI | 0.25409544 | -0.7496409 | -0.3205329 | 0.5694352 | 0.85457899 | 0.21434416 | 0.75523039 | 1 |

Table S5: The number and proportion of 328 and 402 total SNPs identified from the GEA methods, and the number and proportion of genic SNPs for each climate variable.

| | **BIO1** | | | **BIO4** | | | **BIO5** | | | **BIO6** | | |
|---|---|---|---|---|---|---|---|---|---|---|---|---|
| | **Total nSNP** | **Proportion Total SNPs (%)** | | **Total nSNP** | **Proportion Total SNPs (%)** | | **Total nSNP** | **Proportion Total SNPs (%)** | | **Total nSNP** | **Proportion Total SNPs (%)** | |
| *E. blakelyi* | 75 | 22.87 | | 33 | 10.06 | | 38 | 11.59 | | 38 | 11.59 | |
| *E. tereticornis* | 75 | 18.66 | | 130 | 32.34 | | 67 | 16.66 | | 87 | 21.64 | |
| | **Genic nSNP** | **Proportion BIO1 SNPs (%)** | **Proportion Total SNPs (%)** | **Genic nSNP** | **Proportion BIO4 SNPs (%)** | **Proportion Total SNPs (%)** | **Genic nSNP** | **Proportion BIO5 SNPs (%)** | **Proportion Total SNPs (%)** | **Genic nSNP** | **Proportion BIO6 SNPs (%)** | **Proportion Total SNPs (%)** |
| *E. blakelyi* | 28 | 37.33 | 8.54 | 8 | 24.24 | 2.44 | 7 | 18.42 | 2.13 | 5 | 13.16 | 1.52 |
| *E. tereticornis* | 29 | 38.67 | 7.21 | 3 | 2.31 | 0.75 | 5 | 7.46 | 1.24 | 11 | 12.64 | 2.74 |
| | **BIO12** | | | **BIO14** | | | **BIO18** | | | **AAI** | | |
| | **Total nSNP** | **Proportion Total SNPs (%)** | | **Total nSNP** | **Proportion Total SNPs (%)** | | **Total nSNP** | **Proportion Total SNPs (%)** | | **Total nSNP** | **Proportion Total SNPs (%)** | |
| *E. blakelyi* | 24 | 7.32 | | 24 | 7.32 | | 91 | 27.74 | | 75 | 22.87 | |
| *E. tereticornis* | 77 | 19.15 | | 62 | 15.42 | | 99 | 24.63 | | 77 | 19.15 | |
| | **Genic nSNP** | **Proportion BIO12 SNPs (%)** | **Proportion Total SNPs (%)** | **Genic nSNP** | **Proportion BIO14 SNPs (%)** | **Proportion Total SNPs (%)** | **Genic nSNP** | **Proportion BIO18 SNPs (%)** | **Proportion Total SNPs (%)** | **Genic nSNP** | **Proportion AAI SNPs (%)** | **Proportion Total SNPs (%)** |
| *E. blakelyi* | 1 | 4.17 | 0.30 | 6 | 0.25 | 1.83 | 4 | 4.40 | 1.22 | 6 | 8.00 | 1.83 |
| *E. tereticornis* | 5 | 6.49 | 1.24 | 5 | 8.06 | 1.24 | 4 | 4.04 | 1.00 | 8 | 10.39 | 1.99 |

Figure S1: Pairwise climate variation between *E. blakelyi* (blue) and *E. tereticornis* (red). (a) Top panel compares $T_{MA}$ (BIO1) with $P_{MA}$ (BIO12); (b) middle panel compares $T_{Max}$ (BIO5) with $P_{DM}$ (BIO14); and (c) bottom panel compares $T_{Max}$ with $P_{WARMQ}$ (BIO18). Grey circles represent the species' observation records and blue or red points corresponds to sample sites for each species respectively.

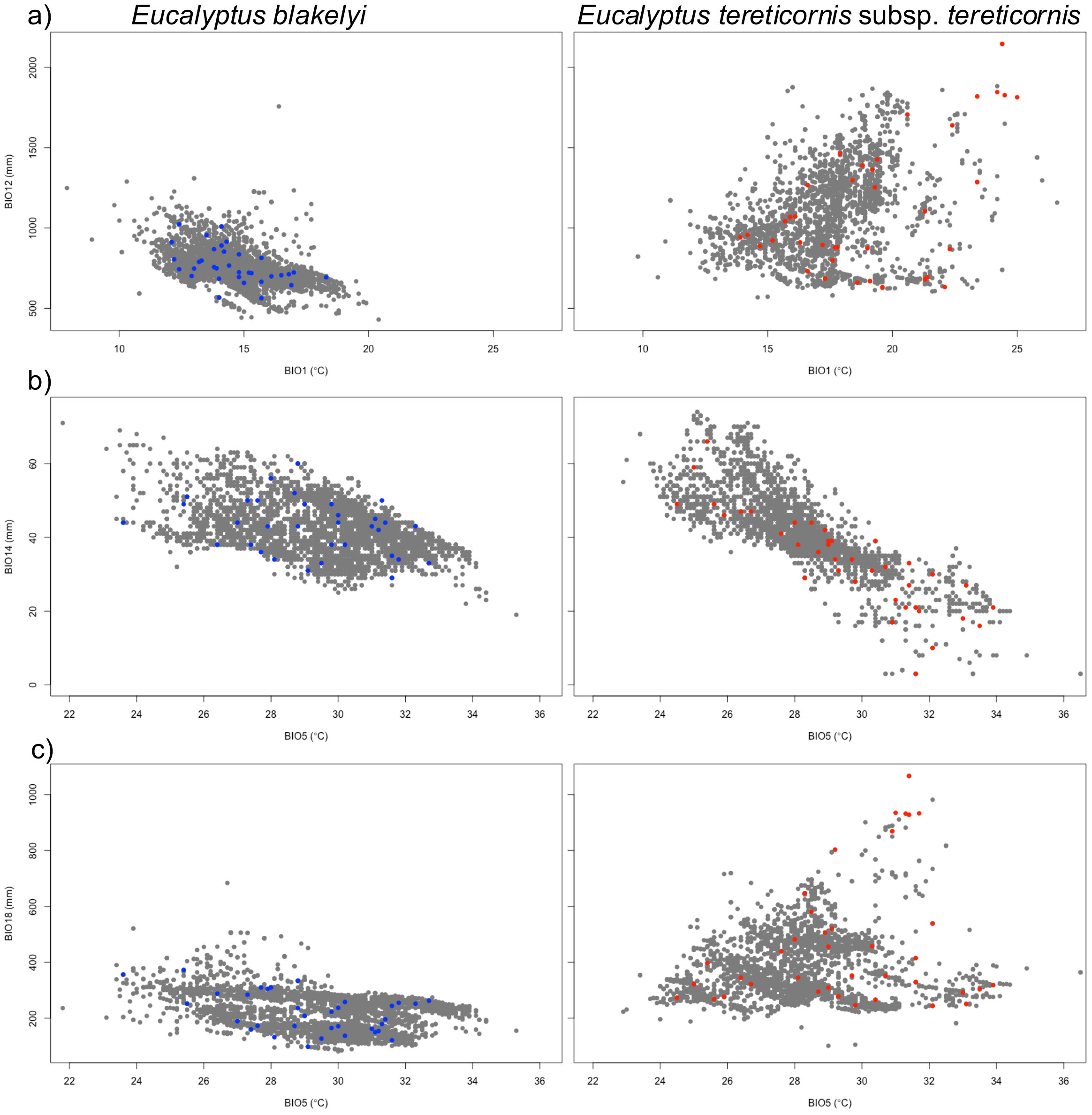

Figure S2: Bayesian Information Criterion (BIC) score plots prior to construction of the Discriminate Analysis of Principal Components plots for (a) *E. blakelyi* and (b) *E. tereticornis.* The lowest BIC score indicates the number of clusters present in the dataset.

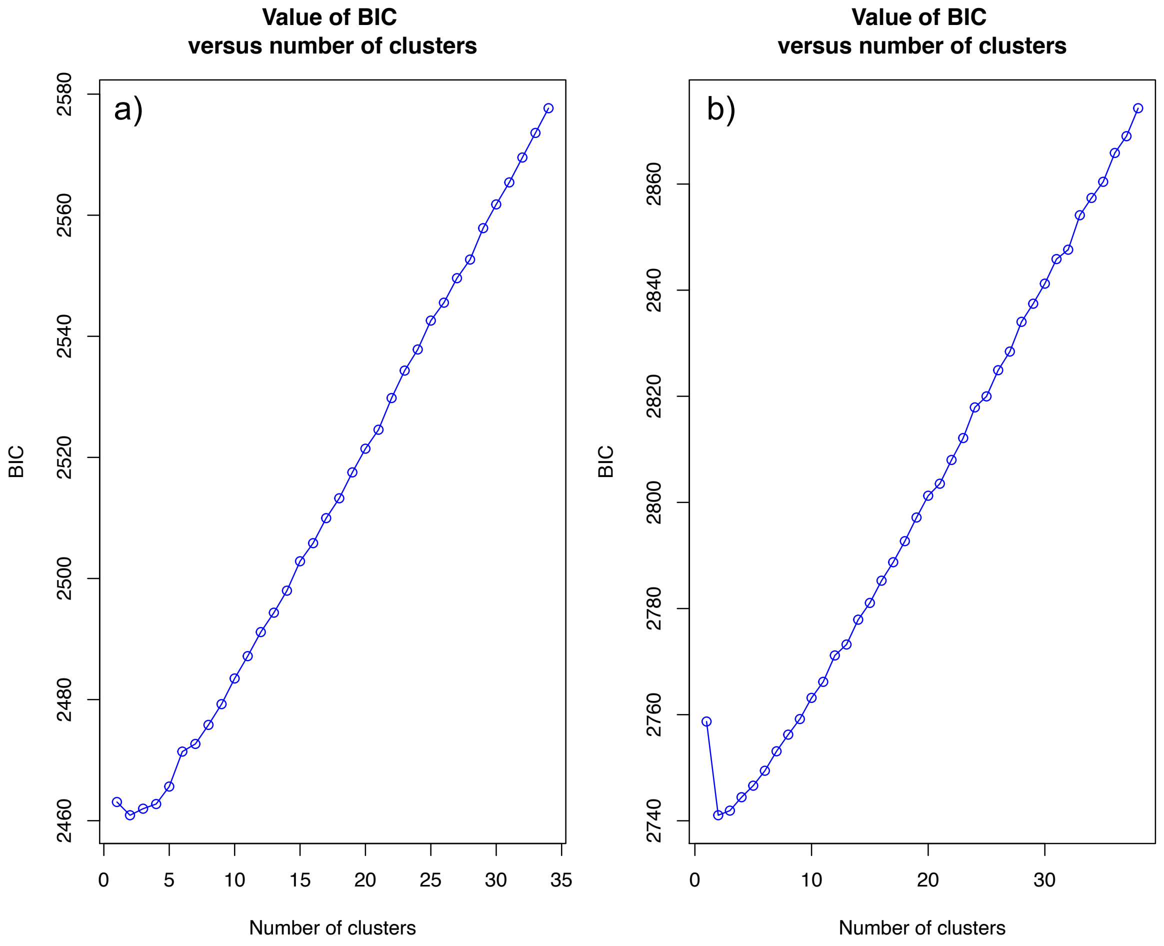

Figure S3: Discriminate analysis of principal components (DAPC) calculated for both *E. blakelyi* and *E. tereticornis*. Number of PCA eigenvalues differ between species; (a) 230 PCs for *E. blakelyi*, and (b) 225 PCs for *E. tereticornis*. For both species, the first two DA eigenvalues were retained. Circles represent 95% confidence intervals, and where circles overlap indicate populations are not significantly different to one another.

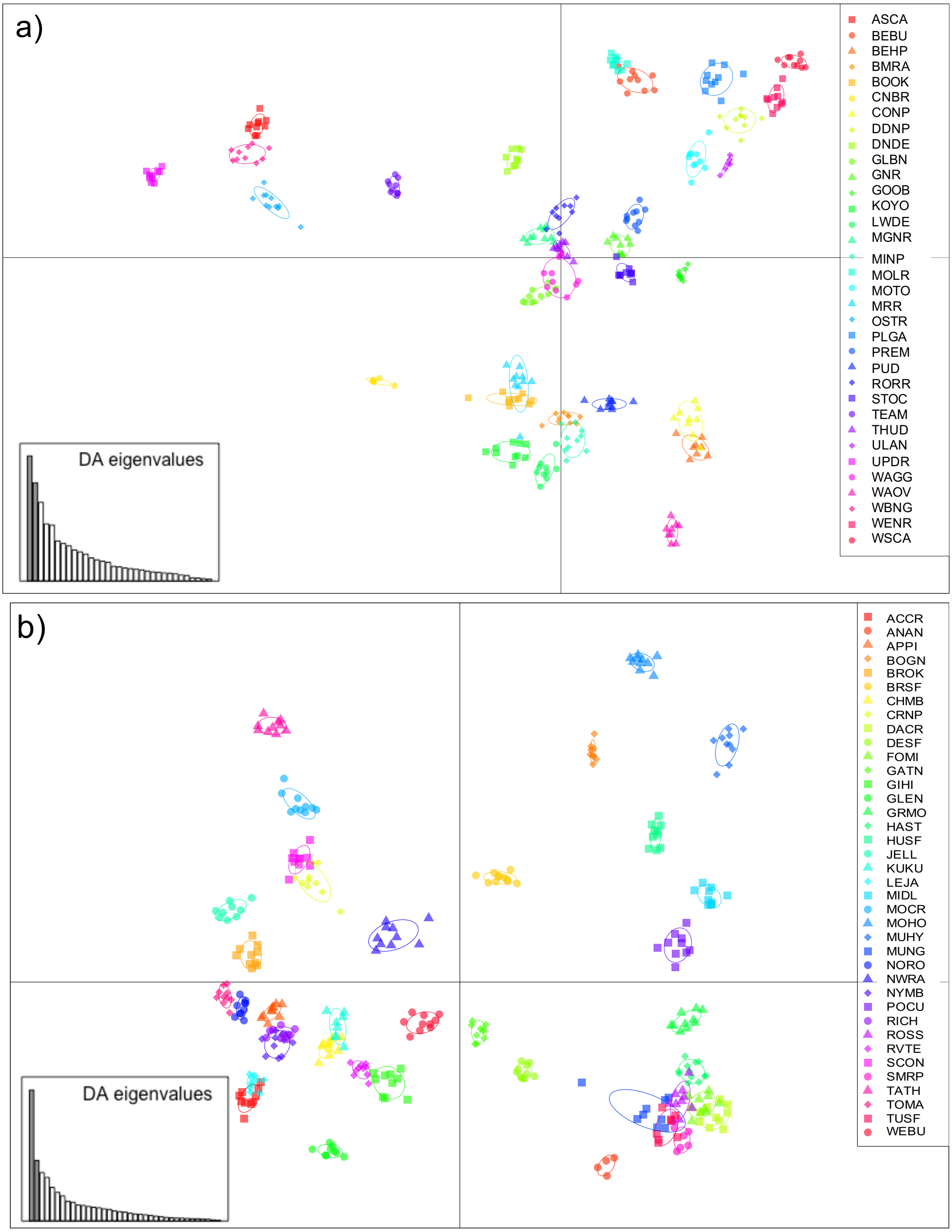

Figure S4: RDA output of the first 2 axes for (a) *E. blakelyi*, and (b) *E. tereticornis* with the eight climatic variables. MAT = mean annual temperature (BIO1); TSN = temperature seasonality (BIO4); MaxT = mean maximum temperature of the warmest month (BIO5); MinT = mean minimum temperature of the coldest month (BIO6); MAP = mean annual precipitation (BIO12); PreDM = mean precipitation of the driest month (BIO14); PreWarmQ = mean precipitation of the warmest quarter (BIO18); AAI = mean annual aridity index.

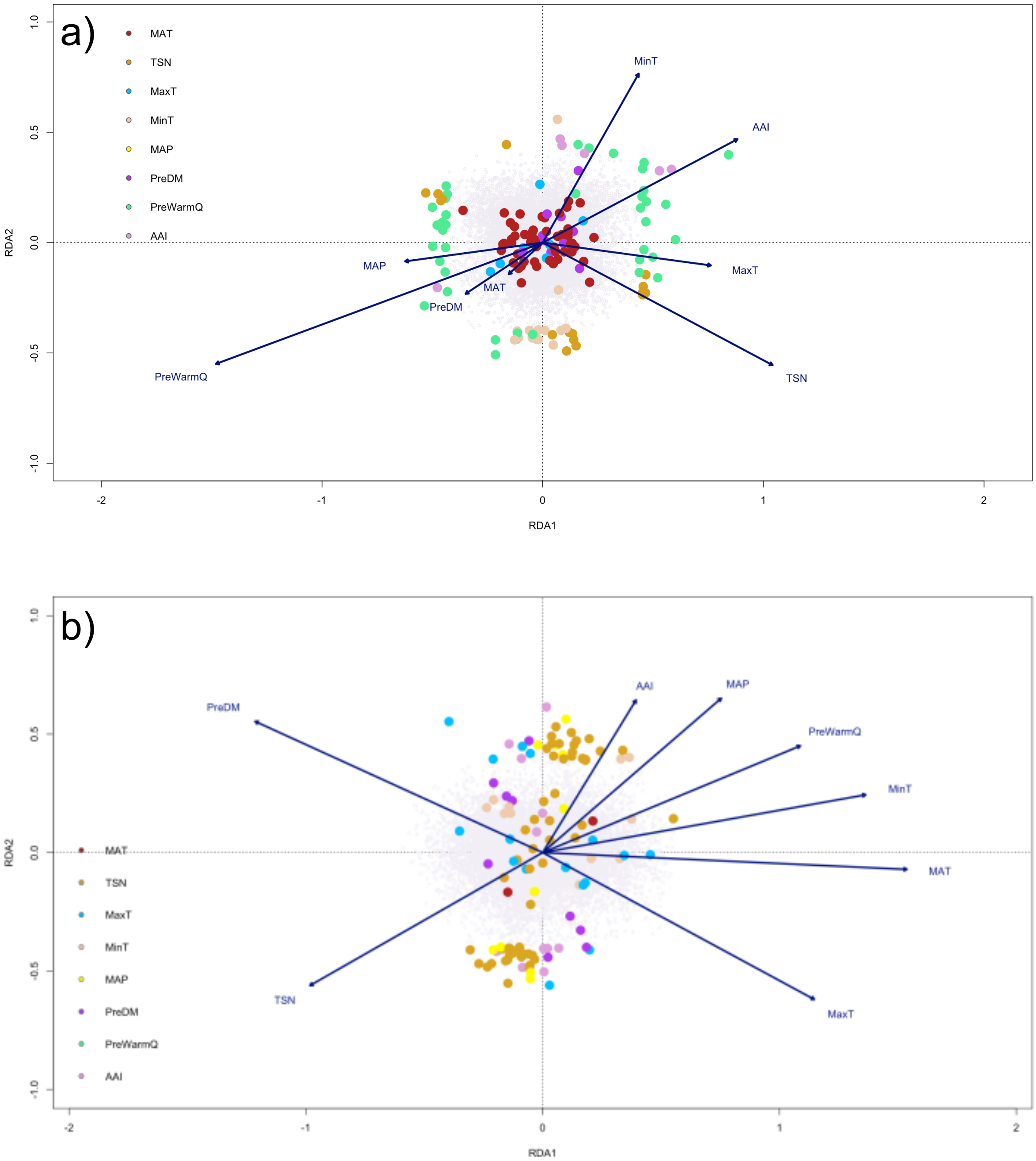
